# A loss-of-function mutation in IL-17F enhances susceptibility of mice to oropharyngeal candidiasis

**DOI:** 10.1101/2020.04.09.028027

**Authors:** Chunsheng Zhou, Leticia Monin, Rachael Gordon, Felix E.Y. Aggor, Rami Bechara, Tara N. Edwards, Daniel H. Kaplan, Sebastien Gingras, Sarah L. Gaffen

## Abstract

Oropharyngeal candidiasis (OPC) is an opportunistic infection of the oral mucosa caused by the commensal fungus *C. albicans*. IL-17 receptor signaling is essential to prevent OPC in mice and humans, but the individual roles of its ligands, IL-17A, IL-17F and IL-17AF, are less clear. A homozygous IL-17F deficiency in mice does not cause OPC susceptibility, whereas mice lacking IL-17A are moderately susceptible. In humans, a rare heterozygous mutation in IL-17F (IL-17F.S65L) was identified that causes chronic mucocutaneous candidiasis, suggesting the existence of essential antifungal pathways mediated by IL-17F and/or IL-17AF. To investigate the role of IL-17F and IL-17AF in more detail, we exploited this ‘experiment of nature’ by creating a mouse line bearing the homologous mutation in IL-17F (Ser65Leu) by CRISPR/Cas9.The resulting *Il17f*^*S65L/S65L*^ mice showed increased susceptibility to OPC, but only in homozygous, not heterozygous, mutant mice. The mutation was linked to impaired CXC chemokine expression and neutrophil recruitment to the infected tongue but not to alterations in antimicrobial peptide expression. These findings suggest mechanisms by which the enigmatic cytokine IL-17F contributes to host defense against fungi.

## Introduction

Fungal infections have a serious impact on human health, yet typically receive less attention than other pathogens (1). *Candida albicans* is a commensal fungus in humans, colonizing mucosal surfaces such as the oral cavity, vaginal tract and gut as well as skin. Although normally benign in healthy individuals, *C. albicans* can cause pathogenic infections such as oropharyngeal candidiasis (OPC, oral thrush) under settings of immunodeficiency. Immunity to *C. albicans* is highly dependent on CD4^+^ T cells, as up 95% of HIV^+^ patients suffer from recurrent OPC during progression to AIDS, correlating with T cell counts (2, 3).

Human T cell responses to *C. albicans* are dominated by Th17 cells (4-7). IL-17A (IL-17) is the signature cytokine of Th17 cells and is also expressed by various subsets of innate lymphocytes such as *γδ*-T, NKT and ILC3 cells (8). IL-17 plays a critical role in antifungal immunity to *C. albicans* (9, 10). Studies in mice demonstrated the importance of IL-17 signaling against *C. albicans* oral infections, as mice lacking IL-17RA, IL-17RC or the signaling adaptor Act1 are all highly sensitive to systemic, oral and dermal candidiasis (11-15). Clinical data from cohorts of chronic mucocutaneous candidiasis disease (CMCD) patients confirmed that susceptibility to candidiasis is strongly associated with IL-17/Th17 deficiencies. Mutations in IL-17RA, IL-17RC or Act1 in humans, as in mice, cause susceptibility to mucocutaneous candidiasis (16-19). Clinical treatment with anti-IL-17A antibodies for autoimmunity is also linked to OPC (20). Similarly, *STAT3, RORC*, and *STAT1* mutations are linked to reduced Th17 frequencies and CMCD (21-23).

The IL-17 family of cytokines is structurally distinct from other cytokine subclasses, composed of IL-17A, IL-17B, IL-17C, IL-17D, IL-17E (IL-25) and IL-17F (24). Among these, IL-17A shares the most homology with IL-17F at the amino acid level (56%) (25, 26). Both IL-17A and IL-17F form homodimers but can also form a heterodimer (IL-17AF) (27-29). All three isoforms signal through the IL-17RA:IL-17RC receptor complex, though the ligands have different binding affinities for the receptor, with IL-17A > IL-17AF > IL-17F (28, 30, 31). Recent data suggest that IL-17A may also signal through an IL17RA:IL-17RD receptor (32), and that IL-17C homodimer may also be competent for signaling (31, 33). Hence our understanding of the IL-17 cytokine family is still developing.

IL-17A and, to a lesser extent, IL-17F activate a program of inflammation in target cells, mainly fibroblasts and epithelial cell types (34, 35). IL-17A induces a characteristic gene signature that includes cytokines (IL-6, GM-CSF, G-CSF), chemokines (CXCL1, CXCL2, CXCL8, CCL2, CCL7, CCL20), matrix metalloproteinases (MMP1, 2, 3, 9, 13), transcriptional and post-transcriptional regulators (C/EBPs, I*κ*B*ξ*, Regnase-1, Arid5a) and antimicrobial peptides (AMPs) (β-defensins, S100A proteins, lipocalin 2) (36-40). Emerging studies also implicate IL-17A in metabolic and proliferation gene expression (41, 42). Though less well characterized, IL-17F induces a similar but not entirely overlapping panel of genes (43-46). Studies of the *in vitro* functions of the IL-17AF heterodimer are comparatively limited, but studies reported to date indicate a similar gene induction profile induced by IL-17AF compared to IL-17A and IL-17F (27, 28).

Despite their capacity to bind the same receptor, IL-17A and IL-17F exert distinct activities *in vivo. Il17a*^*-/-*^ and *Il17f*^*-/-*^ mice show differential susceptibilities to various diseases, both infectious and autoimmune (47, 48). This dichotomy is similarly illustrated in OPC; whereas *Il17a*^*-/-*^ or mice treated with IL-17A-neutralizing antibodies exhibit elevated susceptibility to OPC compared to immunocompetent control mice (49), *Il17f*^*-/-*^ or mice treated with IL-17F neutralizing antibodies are fully resistant to *C. albicans* infection. However, there is evidence that IL-17F does participate in immunity to OPC, as dual blockade of IL-17A and IL-17F increases susceptibility to OPC over blockade of IL-17A alone (49, 50). In line with this, humans with autoimmune polyendocrinopathy candidiasis ectodermal dystrophy (APECED) caused by *AIRE* gene deficiencies have circulating auto-antibodies against both IL-17A and IL-17F, which is thought to underlie susceptibility to CMCD (51-53).

The physiological role of IL-17AF is still unclear. Although IL-17AF can be reliably detected (e.g., by sandwich ELISA), there are no commercial antibodies that block this isoform selectively or efficiently, and hence determining its specific role *in vivo* is challenging. Hints regarding IL-17F and IL-17AF function in came from an unusual cohort of rare CMCD patients discovered by Puel *et al.* that carry a heterozygous serine-to-leucine mutation in the *IL17F* gene at position 65 (IL-17F.S65L) (16). The mutated residue in these individuals lies within the region of interaction of IL-17F with the IL-17 receptor. *In vitro*, this mutation impaired IL-17F binding to the receptor but had no apparent impact on the ability of IL-17F to dimerize with IL-17F or IL-17A. Cultured human fibroblasts treated with an IL-17F homodimer containing this mutation (IL-17FS65L/IL-17F or IL-17S65L/IL-17FS65L) or a mutant IL-17AF heterodimer (IL-17A/IL-17FS65L) showed strongly impaired signaling *in vitro* (16). Accordingly, the S65L mutation in human IL-17F appeared to be both a loss of function and a dominant negative mutation, causing functional blockade of both IL-17F and IL-17AF. Importantly, IL-17A homodimers were still found in these patients yet were apparently insufficient to fully protect from candidiasis. These data thus implied that IL-17F and/or IL-17AF are significant contributors to the antifungal immune response in humans.

In this study, we created an IL-17F.S65L mouse strain using CRISPR/Cas9 technology to exploit this “experiment of nature” (16) in order to better understand the functions of IL-17F and IL-17AF in immune responses. We found that *Il17f*^*S65L/S65L*^ mice exhibited a similar susceptibility to OPC as *Il17a*^*-/-*^ mice, which contrasted with the known resistance of *Il17f*^*-/-*^ mice to OPC (49). There was no detectable disease susceptibility in mice heterozygous for the mutation, potentially suggesting differences between mouse and human IL-17F function *in vivo.* The increased susceptibility of these mice to fungal infection was linked to impaired expression of CXC chemokines and concomitantly reduced neutrophil recruitment to the oral mucosa, but surprisingly not to expression of key antimicrobial peptides known to control OPC such as *β*-defensin-3.

## Methods

### Generation of Il17f^S65L/S65L^ knockin mice

*Il17f*^*S65L/S65L*^ mice were created by CRISPR/Cas9 by the Transgenic and Gene Targeting (TGT) and Innovative Technologies Development (ITD) Core facilities in the Department of Immunology, University of Pittsburgh. Briefly, a S.py. Cas9 target sequence overlapping the Ser65 codon in the mature *Il17f* sequence (following signal peptide cleavage) was selected: GTTCCCCTCAGAGATCGCTG **AGG**. The protospacer adjacent motif (PAM) in bold was not included in the sgRNA. Cas9 mRNA and the sgRNA were produced as described (54, 55). A 127-mer oligonucleotide (Ultramer, IDT) was used as template for homology-directed repair (HDR): *Il17f*-S65L-HDRv3: 5’-CATCCTGCTTTACTTTTTATTTTTTTCCTTCAGCATCACTCGAGACCCCCACCGGTTCC C<u>TCTAG</u>AAATCGCTGAGGCCCAGTGCAGACACTCAGGCTGCATCAATGCCCAGGGT CAGGAAGACAGC-3’. The oligonucleotide contains a substitution to convert Serine 65 to Leucine, which contemporaneously disrupts an HPY188I restriction site (underlined) to facilitate genotyping. The sequence also contains an additional silent substitution, bringing a total of 4 mismatches (red, above) between the sgRNA target sequence and the designed allele, thus limiting further re-editing of the mutant allele by Cas9. C57BL/6J embryos were microinjected with the sgRNA (50 ng/µl), the HDR oligonucleotide (0.5 µM) and Cas9 mRNA (100 ng/µl). Embryos that developed to the 2-cell stage were transferred into pseudopregnant female surrogates. Sixteen founders were identified by PCR amplification of the target region (Forward: 3’-ATGGGAGAAACCCCGTTTTA-5’; reverse: 3’-TCCAACCTGAAGGAATTAGAACA-5’) followed by restriction digestion of the PCR product with HPY188I. The correct sequence was validated by Sanger sequencing of the PCR products following TOPO cloning. Of these, 5 founders were homozygous for the mutation (**Supp. Table 2**). Based on the CRISPOR website (56), there were only 2 potential off-target sequences with fewer than 4 mismatches in the mouse genome (**Table 1**). Among all founders analyzed, there were no mutations introduced at those sites, as determined by PCR and Sanger sequencing. The expanded line was backcrossed twice to C57BL/6J mice prior to colony expansion.

**Table 1.**
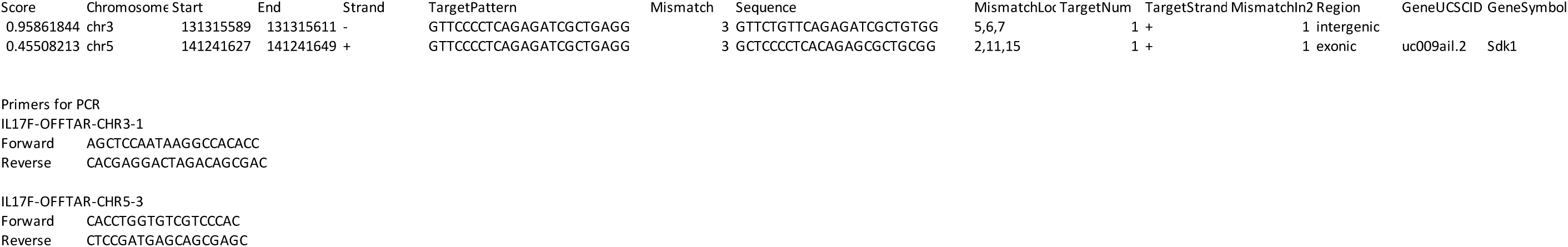
Off target screening of Il17f S65L mutant mice. Two off-target sites with up to 3 bp mismatches were predicted in creation of *Il17f*^*S65L/S65L*^ mice, located on chromosomes 3 and 5. Precise locations and sequences are shown. PCR primers used to amplify and sequence these regions are indicated. See Methods section for details.

### Other mice

*Il17f*^*Thy1.1*^ reporter mice were previously described (57). *Il17ra*^-/-^ mice were a gift from Amgen. *Act1*^*-/-*^ mice were from U. Siebenlist, NIH. WT mice were from JAX, Taconic Farms, or generated from breeding colonies. All mice were on the C57BL/6 background. Age matched mice (6-10 weeks) were used for experiments with both sexes. Animal use protocols were approved by the University of Pittsburgh Institutional Animal Care and Use Committee.

### Model of oropharyngeal candidiasis (OPC)

OPC was induced by sublingual inoculation with a cotton ball saturated in *C. albicans* (CAF2-1) for 75 min under anesthesia, as described (58). Tongue homogenates were prepared on a gentleMACS (Miltenyi Biotec), and fungal burdens determined by plating on YPD agar with Ampicillin. Anti-IL-17A or isotype control antibodies (Bio X Cell) (200ug/ul) were administered i.p. on days -1, 1, and 3 relative to *C. albicans* infection.

### Flow cytometry

Tongues were digested with Collagenase IV (0.7mg/ml) in HBSS for 30 min. Cell suspensions were separated by Percoll gradient centrifugation. Abs were from eBioscience, BD Biosciences, and BioLegend (59). Data were acquired on an LSR Fortessa and analyzed with FlowJo (Ashland, OR).

### qPCR

Tongues were homogenized in in a Gentle MACS Dissociator (Miltenyi Biotec) and tongue RNA was extracted using RNeasy kits (Qiagen). RNA from ST2 cells was extracted in RLT buffer. cDNA was generated using a SuperScript III First Strand Synthesis System (Invitrogen). Relative quantification of gene expression normalized to *Gapdh* was determined by real-time PCR with SYBR green (Quanta BioSciences) on the Applied Biosystems 7500 platform. Primers were from QuantiTect (Qiagen).

### Cell culture and Th17 differentiation

The ST2 stromal cell line was cultured in *α*-minimum essential medium (α-MEM; Sigma-Aldrich, St. Louis MO) with L-glutamine, antibiotics, and 10% fetal bovine serum. 5×10^%^ cells were seeded into 6 wells plates prior to cytokine stimulation. Recombinant IL-17F.S65L and IL-17F were synthesized by Bon Opus Biosciences (Millburn, NJ) by expression in Expi293 cells (ThermoFisher). TNF*α* was from Peprotech and used at 2 ng/ml (Rocky Hill, NJ).

Naïve splenic CD4^+^ T cells were purified by magnetic separation (Miltenyi Biotec). T cells were activated by α-CD28 (5 ug/ml; BioXCell) and plate-bound α-CD3 (clone 145-TC11, 5 ug/ml; BioXCell) in complete RPMI medium supplemented with 10% fetal bovine serum, 2 mM L-glutamine, 100 U/ml penicillin, 100 mg/ml streptomycin, 50 mM 2-*β*-mercaptoethanol, and sodium pyruvate) for 4 d with IL-1β (50 ng/ml), IL-6 (50 ng/ml), IL-23 (50 ng/ml), and TGFβ (5 ng/ml). Cytokines were from R&D Systems.

### Statistics

Data were analyzed on Prism (Graphpad) using ANOVA or Student’s t test. Fungal burdens are presented as geometric mean analyzed by ANOVA with Mann-Whitney analysis. \**P* < 0.05; **< 0.01; *** < 0.001; **** < 0.0001.

## Results

### Murine IL-17F S65L is a loss-of-function mutation

The S65L residue in IL-17F that is mutated in humans with CMCD is conserved among many species, including mice (16). The human IL-17F.S65L cytokine has no signaling capacity *in vitro* (16). To determine whether the mouse orthologue of IL-17F.S65L functions similarly to its human counterpart, a His-tagged murine IL-17F.S65L and a corresponding wild type IL-17F control were expressed in Expi293 cells and purified from conditioned media on a nickel-charged affinity resin (Ni-NTA) (**Fig. 1A**). Recombinant IL-17F.S65L and IL-17F migrated according to their predicted dimeric sizes on non-reducing SDS-PAGE (**Fig. 1B**). The multiple bands in the reduced (R) and non-reduced (NR) gels are consistent with the expectation that these recombinant cytokines exist in multiple glycosylated and non-glycosylated forms (**Fig. 1B, Suppl. Fig. 1**).

**Figure 1.**
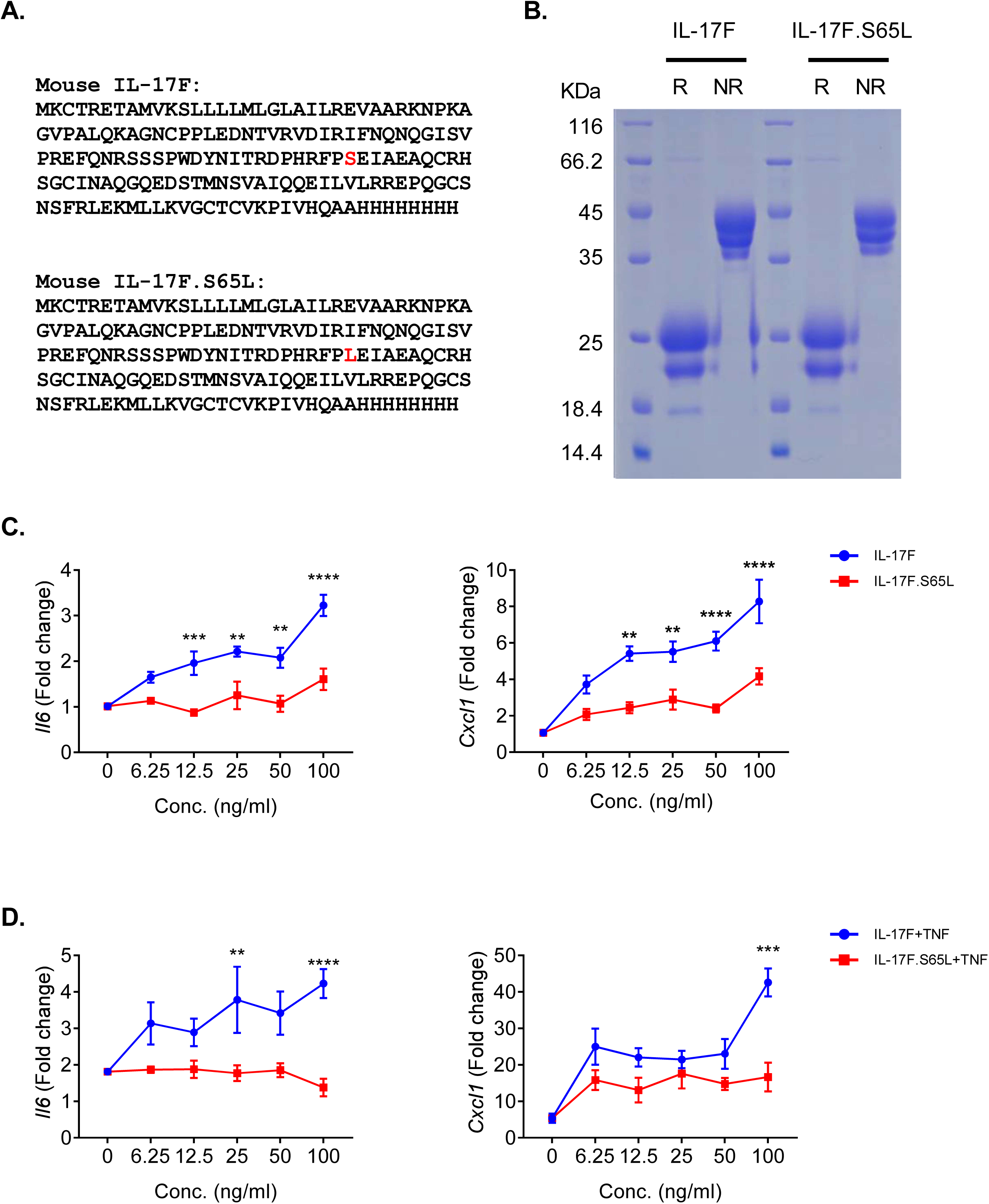
Murine IL-17F.S65L is a loss-of-function mutation. (A) Amino acid sequences of recombinant His-tagged WT IL-17F and the IL-17F.S65L mutant used in this study. The Ser65 residue is shown in red. (B) IL-17F and IL-17F.S65L were transfected into Expi293 cells. Culture medium was collected on day 5 post transfection purified by Ni-NTA. The eluted fraction was analyzed by SDS-PAGE in reducing (R) condition and non-reducing (NR) conditions. Size markers are indicated at left. (C, D) ST2 stromal cells were treated with the IL-17F (blue) and IL-17F.S65L (red) at increasing doses in the absence (C) or presence (D) of TNF*α* (2 ng/ml) for 3 h. *Il6* or *Cxcl1* mRNA was assessed by qPCR. Data is graphed as fold-change relative to untreated conditions ± SEM from 2 independent experiments, analyzed by ANOVA. \*\**P* < 0.01, *** < 0.001, **** < 0.0001.

To evaluate the signaling capability of murine IL-17F.S65L, murine ST2 stromal cells were treated with increasing concentrations of IL-17F.S65L or WT control IL-17F for 3 h. Two IL-17F-inducible genes, *Il6* and *Cxcl1*, were assessed by qPCR as endpoints of signaling. As expected, control IL-17F induced *Il6* and *Cxcl1* in a dose-dependent manner (**Fig. 1C**). In contrast, IL-17F.S65L at most doses (6.25-50 ng/ml) did not detectably upregulate *Il6* or *Cxcl1*. However, both mRNAs were slightly enhanced by IL-17F.S65L at a supraphysiological dose (100 ng/ml), but nonetheless showed significantly reduced activity compared to non-mutated IL-17F (**Fig. 1C**). Because IL-17F induces gene expression synergistically with other cytokines (43, 46), we evaluated IL-17F.S65L responsiveness in concert with a suboptimal dose of TNF*α* (2 ng/ml). Control IL-17F synergized with TNF*α* to induce *Il6* and *Cxcl1*, but the IL-17F.S65L mutant did not synergistically enhance expression of these genes (**Fig. 1D**). Thus, analogous to the IL-17F.S65L mutation found in CMCD patients, a mouse IL-17F.S65L homodimer appears to be a loss-of-function mutation.

### Il17f^S65L/S65L^ mice are modestly susceptible to OPC

To determine the function of the murine IL-17F.S65L mutation *in vivo*, we created an IL-17F.S65L mutant mouse strain by CRISPR/Cas9 technology. Sixteen founders were generated, of which 5 had a homozygous nucleotide replacement (**Fig. 2A, Suppl. Fig. 2**). None of the founder lines were found to have mutations in the either of the two predicted off-target sites, ascertained by PCR of genomic DNA and sequencing (**Table 1**, see Methods section for details). Similar to *IL17a*^*-/-*^ and *Il17f*^*-/-*^ mice (47), there were no obvious abnormalities in the health of these mice when maintained in SPF conditions, and they bred normally with Mendelian numbers of offspring.

**Figure 2.**
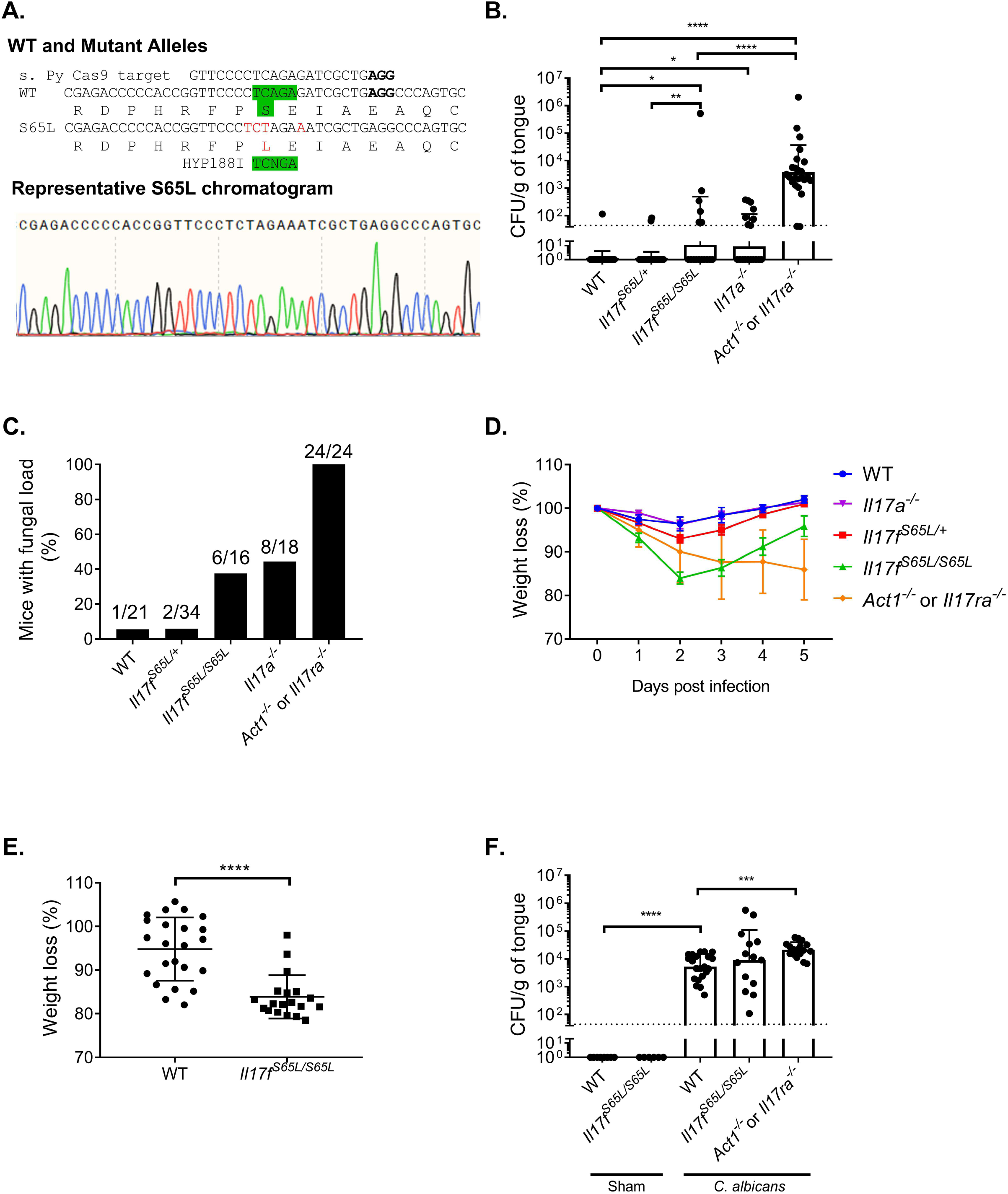
Il17f^S65L/S65L^ mice are modestly susceptible to OPC. (A) Top: Schematic diagram of murine IL-17F.S65L substitution created by CRISPR/Cas9 including disruption of an endogenous HPY188I restriction site. Bottom: Representative chromatogram of genomic DNA sequencing from a representative founder mouse. (B) The indicated mice were infected sublingually with *C. albicans* or PBS (Sham). Fungal burdens were assessed by CFU enumeration on YPD/Amp on day 5 p.i. Graphs show geometric mean + SD. Dashed line indicates limit of detection (∼30 CFU/g). Data were compiled from five 5 experiments. Each symbol represents one mouse. (C) Data from panel B is represented as percentage of mice with detectable fungal load in the tongue. Values above indicate number of mice with fungal burden/total. (D) Weight in the animals from panel B was assessed daily and percentage loss relative to day 0 is shown for each time point. (E) Weight loss of *Il17f*^*S65L/S65L*^ and WT mice at day 2 p.i. from panel B. Graph shows mean ± SD. (F) Fungal burdens were assessed on day 2 p.i.. Data are compiled from 3 independent experiments. Data were analyzed by ANONVA or Student’s t-test, with Mann-Whitney analysis for fungal load analysis. \**P* < 0.05, ** < 0.01, *** < 0.001, **** < 0.0001.

Since humans with this mutation experience CMCD as heterozygotes (no homozygotes were described (16)), we hypothesized that mice that are both homozygous and heterozygous for the IL17F.S65L mutation would be susceptible to mucosal candidiasis. Accordingly, mice were subjected to OPC by a 75 minute sublingual exposure to *C. albicans* (strain CAF2-1) by standard methods (58, 60). In this model, WT mice typically clear *C. albicans* from the oral mucosa within 3 days and show no overt signs of illness such as prolonged weight loss (12, 61). We previously demonstrated that mice lacking IL-17RA or Act1 maintain a high fungal burden in the oral cavity after infection, which we typically measure at day 5 post infection (p.i.) (12, 14), Mice lacking IL-17A (*Il17a*^*-/-*^ or given anti-IL-17A antibodies) have detectable fungal burdens, though consistently lower than *Il17ra*^*-/-*^ or *Act1*^*-/-*^ mice (49). In contrast, mice lacking IL-17F, either by knockout or with neutralizing antibodies, are fully resistant to OPC in this system (49). Here, we observed that *Il17f*^*S65L/S65L*^ mice had a significantly higher oral fungal burden compared to WT controls at 5 days p.i., at levels similar to *Il17a*^*-/-*^ animals (**Fig. 2B**). Notably, however, the *Il17f*^*+/S65L*^ heterozygous mice did not have elevated fungal loads compared to WT. Approximately 40% of the *Il17f*^*S65L/S65L*^ and *Il17a*^*-/-*^ mice still had a detectable fungal load at this time point, whereas the WT and the *Il17*^*/S65L/+*^ heterozygous mice almost all fully cleared the infection (**Fig. 2C**). *Il17f*^*S65L/S65L*^ mice also lost slightly more weight than WT controls, which was most evident at day 2 p.i. (**Fig. 2D, E**). However, oral fungal burdens in *Il17f*^*S65L/S65L*^ mice were not measurably different at day 2 (**Fig. 2F**). Collectively, these results indicate that IL-17F.S65L mutation contributes detectably, albeit modestly, to susceptibility to OPC, but only when the mutation is present on both alleles.

### Il17f^S65L/S65L^ mice have impaired neutrophil recruitment during OPC

To understand the immunological mechanisms by which the IL-17F.S65L mutation promotes susceptibility to OPC, we evaluated factors known to be critical for antifungal immunity mediated by IL-17R signaling (12, 15, 61). Neutrophils are vital for fungal clearance in OPC (62-64). We have observed that *Il17ra*^*-/-*^ mice show impaired recruitment of neutrophils to the oral mucosa following OPC induction (12, 15, 65). IL-17F upregulates expression of neutrophil-attracting chemokines such as CXCL1 and CXCL2 (66-68), and both human and murine IL-17FS65L showed impaired induction of this chemokine *in vitro* (**Fig. 1**, (16)). Consistent with this, *Cxcl1* gene expression in the tongue was downregulated in *Il17f*^*S65L/S65L*^ mice upon *C. albicans* oral infection. However, expression of *Cxcl2* was similar between *Il17f*^*S65L/S65L*^ mice and WT control (**Fig. 3A**). Flow cytometry analysis revealed that early neutrophil recruitment to the tongue measured at day 2 p.i. was significantly, though not completely, decreased in infected *Il17f*^*S65L/S65L*^ mice compared to WT (45% versus 57%) (**Fig. 3B**). Thus, impaired neutrophil recruitment in *Il17f*^*S65/S65L*^ mice may be one cause for OPC susceptibility in these mice.

**Figure 3.**
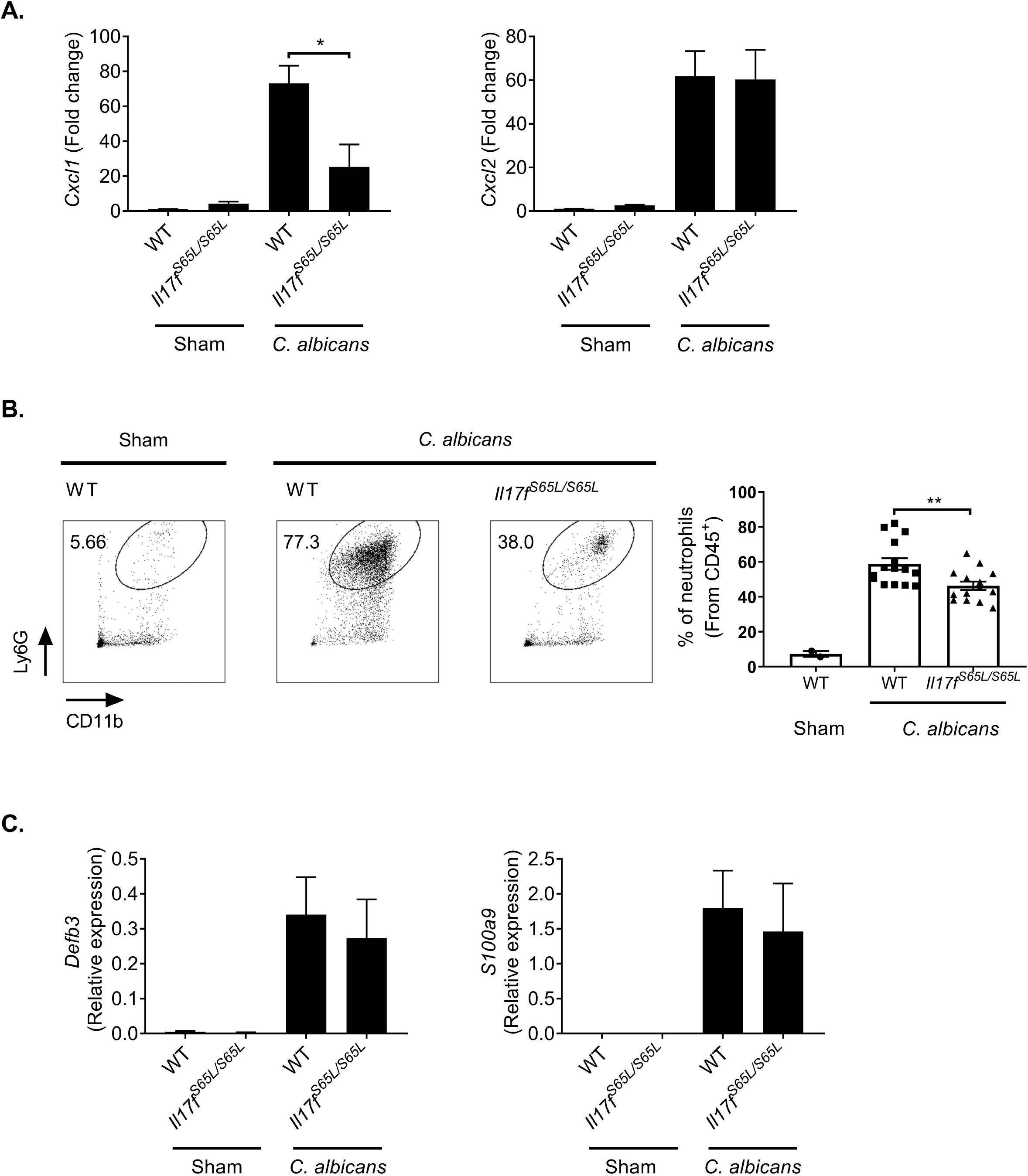
IL-17F.S65L mice exhibit impaired neutrophil recruitment during OPC. The indicated mice were infected sublingually with *C. albicans* or PBS (Sham). (A) Tongues were harvested at day 2 p.i.. Total mRNA from tongue homogenates was assessed by qPCR, normalized to *Gapdh.* Fold change compared to Sham is indicated. Graphs show mean + SEM. Data were merged from 2 independent experiments. (B) Single cell suspensions from tongues harvested at day 2 p.i. were analyzed by flow cytometry. Cells were gated on the CD45^+^ live cell population and stained for CD11b and Ly6G. Left: representative FACS plots. Right: compiled results from 3 independent experiments. Graph shows mean + SEM. (C) Total mRNA from tongue homogenates was assessed for *Defb3* (encoding *β* defensin 3) and *S100a9* by qPCR, normalized to *Gapdh.* Relative expression data are graphed as mean + SEM merged from 4 independent experiments. Data were analyzed by ANOVA. \**P* < 0.05, **P< 0.01.

AMPs such as β-defensins and calprotectin (S100A8/S100A9) have antifungal activity towards *C. albicans* (69-74). IL-17RA knockout mice show impaired AMP expression after *C. albicans* oral challenge (12, 61). Surprisingly, the induction of *Defb3* and *S100a9* in the oral mucosa of *Il17f*^*S65L/S65L*^ mice was equivalent to WT upon oral *C. albicans* infection (**Fig. 3C**). Therefore, the increased disease susceptibility of OPC in *Il17f*^*S65L/S65L*^ mice is apparently not due to inefficient AMP expression. Moreover, these data suggest that IL-17F and/or IL-17AF signaling may be more critical for the neutrophil response than for induction of AMPs.

### Increased OPC in Il17f^S65L/S65L^ mice is not due to impaired IL-17A

IL-17A has an antifungal activity in OPC (49, 75). To determine if IL-17A expression was impacted in *Il17f*^*S65L/S65L*^ mice, we analyzed *Il17a* mRNA from tongue at day 2 p.i. Interestingly, *Il17f*^*S65L/S65L*^ mice showed elevated *Il17a* as well as *Il17f* (**Fig. 4A**), arguing that the higher fungal susceptibility in *Il17f*^*S65L/S65L*^ mice is likely not due to impaired *Il17a* expression; rather, the mildness of the disease susceptibility in these mice could be due to compensatory IL-17A levels. To determine whether the increased *Il17a* expression is directly caused by the IL-17F.S65L mutation or a result of fungus infection in the oral mucosa, we subjected splenic CD4^+^ T cells to *in vitro* differentiation for 3 days under Th17 conditions (IL-6, IL-23, TGFβ). IL-17A concentrations in supernatant were measured by ELISA. As shown, there was no significant difference in the amount of IL-17A produced by T cells obtained from *Il17f*^*S65L/S65L*^, *Il17f*^*-/-*^ or WT mice (**Fig. 4B**). Thus, IL-17F.S65L does not directly influence IL-17A production from T cells.

**Figure 4.**
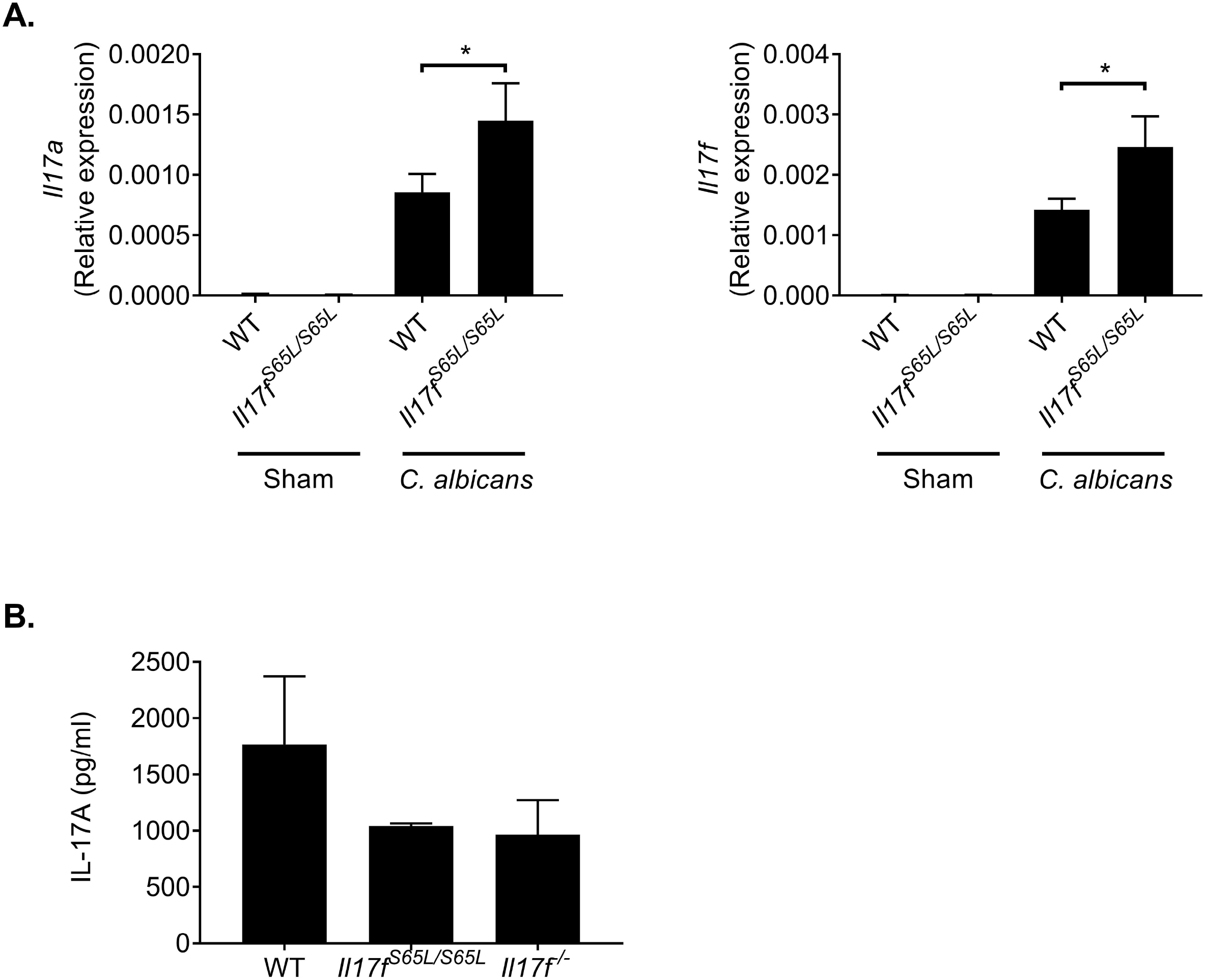
The IL-17F.S65L mutation does not influence IL-17A production from T cells. The indicated mice were infected sublingually with *C. albicans* or PBS (Sham). (A) At 2 days p.i., total mRNA was isolated from tongue and *Il17a* and *Il17f* measured by qPCR and normalized to *Gapdh*. Data show mean + SEM from 4 independent experiments. (B) Naïve CD4^+^ T cells from indicated mice were isolated from spleen and activated with anti-CD3 and anti-CD28 antibodies (5 ug/ml) under Th17 conditions (IL-1β, IL-6, IL-23 at 50 ng/ml and TGFβ at 5 ng/ml) for 4 days. IL-17A in supernatants was measured by ELISA. Data were analyzed by ANOVA. \**P* < 0.05.

To determine if the elevated IL-17A in the tongue could compensate for IL-17F signaling in *Il17f*^*S65L/S65L*^ mice, we treated mice with neutralizing antibodies against IL-17A (75) during OPC induction. In WT mice, blockade of IL-17A caused increased fungal burdens and an increased percentage of mice with fungal loads, measured at day 5 p.i., as previously demonstrated (**Fig. 5A, B**) (49). However, IL-17A blockade in *Il17f*^*S65L/S65L*^ mice did not further increase the oral fungal load compared to *Il17f*^*S65L/S65L*^ isotype control mice or WT mice treated with α-IL-17A antibodies (**Fig. 5A, B**). Thus, in contrast to findings in *Il17f*^*-/-*^ mice (49), IL-17A does not compensate for the IL-17F.S65L mutation during OPC.

**Figure 5.**
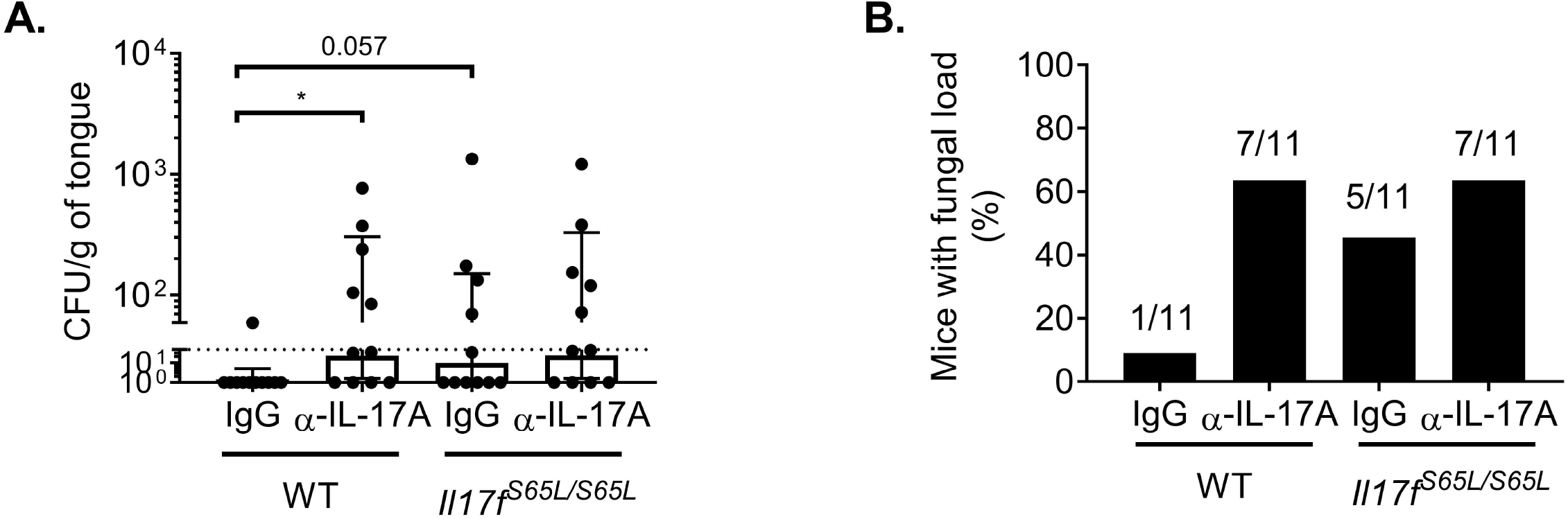
IL-17A does not compensate for IL-17F signaling in Il17f^S65L/S65L^ mice during OPC. The indicated mice were infected sublingually with *C. albicans* or PBS (Sham). (A) Mice were injected i.p. with α-IL-17A neutralizing Abs or isotype control IgG (200ug) on days -1, 1, and 3 relative to the infection. CFU was determined on day 5 p.i.. Data were compiled from 2 independent experiments. (B) Data from panel A is represented as percentage of mice with detectable fungal loads. Values above indicate number of mice with fungal burden/total. \**P* ˂ 0.05 by ANOVA and Mann-Whitney analysis.

### IL-17F is produced dominantly by γd T cells

Unlike humans, mice do not harbor *C. albicans* as a commensal microbe. Multiple studies have verified that the initial response to oral infection with this organism during this first encounter in mice derive entirely from the innate immune compartment (50, 59, 76-79). The sources of IL-17A during acute OPC were previously shown to be dominantly from an unconventional, innate-acting TCRαβ^+^ cell population and *γδ*-T cells (59, 76). Group 3 innate lymphoid cells (ILC3s) were reported to produce IL-17A as well (50). Using *Il17f*^*Thy1.1*^ reporter mice (57) we observed an increased level of *Il17f*-expressing cells 2 days after *C. albicans* infection (**Fig. 6A**), the time point at which *Il17f* mRNA expression peaks (49). *γδ*-T cells constituted the major Thy1.1^+^ population (64%), and TCRβ^+^ cells also comprised a significant portion of the Thy1.1^+^ cells (21%). A population of TCR*γδ*-negative TCR*β*-negative cells (15.5%) that may be ILC3s was also observed (**Fig. 6B**).

**Figure 6.**
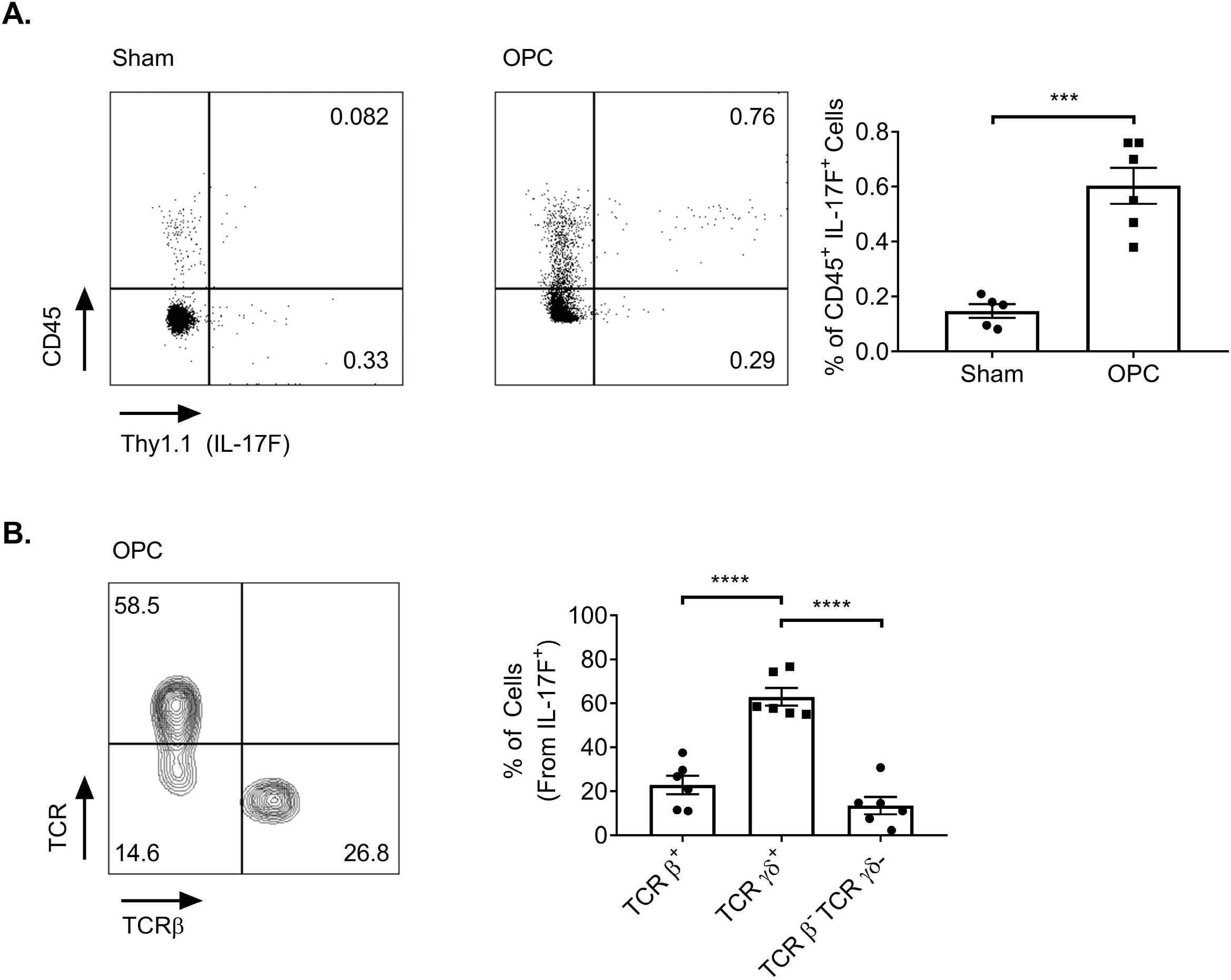
IL-17F is dominantly produced by oral γd-T cells during OPC. *Il17f*^***Thy1.1***^ reporter mice were subjected to OPC. At day 2 p.i., single cell suspensions from tongue were analyzed by flow cytometry. (A) Cells were stained for Thy1.1 (a reporter for IL-17F) and CD45 in the live lymphocyte gate. Left: representative FACS plots. Right: compiled results from 2 independent experiments. (B) Single cell suspensions from panel A were stained for TCRβ and TCRγd in the CD45^+^ Thy1.1^+^ gate. Left: representative FACS plots. Right: compiled results from 2 independent experiments. Data were analyzed by ANOVA or Student’s t-test. \*\*\**P* < 0.001, **** < 0.0001.

## Discussion

*C. albicans* asymptomatically colonizes most healthy individuals and usually only causes mucocutaneous infections in immunocompromised individuals (2, 80). Deficits in IL-17 signaling or Th17 cell development are particularly linked to superficial *C. albicans* infections (21, 81). Although CMCD patients with null mutations in genes encoding IL-17RA, IL-17RC or the adaptor ACT1 have all been described, thus far no CMCD patients have been reported to have a single IL-17A deficiency (22). Even anti-IL-17A biologic drugs used in psoriasis and other autoimmune disease cause only a modest increase in mucocutaneous *Candida* infections (20, 82).

Observations in mouse studies of OPC predicted a role for IL-17 signaling in oral candidiasis (12, 15, 83, 84). Mice treated with IL-17A neutralizing antibodies have lower fungal burdens compared to the *Il17ra*^*-/-*^ mice during OPC (49, 75), analogous to the surprisingly low percentage of patients who only experience mild *C. albicans* mucocutaneous infections during secukinumab treatment (20, 82, 85). Even so, OPC susceptibility in mice does not perfectly phenocopy the human condition. Unlike nearly all humans (5, 86), laboratory mice do not harbor *C. albicans* as a commensal organism, and hence the acute OPC model system mainly reflects events in the innate response (59, 76, 78, 87). Mice lacking STAT3 in T cells are not susceptible to OPC (76). Mice also express different AMP proteins than humans, especially in saliva (88, 89). Of relevance to C. albicans infections, the AMP *β*-defensin 3 is essential to prevent OPC in mice (61), but has no direct orthologue in humans (90, 91). Moreover, the composition of the microbiome varies among species, including in the mouth (92, 93).

Although humans with the IL-17F.S65L mutation are rare, this *Il17f*^*S65L/S65L*^ knockin strain reveals some distinctions between mice and humans. Puel *et al.* reported that 70% of the individuals carrying the IL-17F.S65L mutation had a confirmed diagnosis of mild CMCD (16). In contrast, almost all the *Il17f*^*S65L/+*^ mice fully cleared the fungus in our studies, indistinguishable from WT controls. Rather, susceptibility was only seen in mice carrying the mutation on both alleles, which may argue against a dominant negative role for this mutation in the mouse. Even so, the *Il17f*^*S65L/S65L*^ mice showed fungal susceptibility that was more similar to *Il17a*^*-/-*^ mice than to *Il17f*^*-/-*^ mice, as the latter but not the former are resistant to OPC (49). Although IL-17F-deficient mice were shown to be fully resistant to OPC, it has been shown that blocking both IL-17A together with IL-17F increases susceptibility compared to blockade of IL-17A alone (49, 50). This may be reminiscent of APECED (*AIRE* deficiency) where patients show neutralizing antibodies against multiple Th17 cytokines, including IL-17A and IL-17F (52, 53, 94). However, administration of IL-17A neutralizing antibodies (which also efficiently block IL-17AF (49)) in *Il17f*^*S65L/S65L*^ mice did not further increase susceptibility to OPC, suggesting that any residual IL-17A homodimers present in these animals do not provide additional detectable protection in this model system.

Taken together, the above results are consistent with a protective role for the heterodimer IL-17AF in OPC. Studies in human fibroblasts showed that the IL-17F.S65L mutation impairs signaling of both the IL-17F homodimer and the IL-17AF heterodimer (16). The increased susceptibility to OPC in *Il17f*^*S65L/S65L*^ mice could be due either to a contribution of IL-17F homodimer and/or the IL-17AF heterodimer. Nonetheless, anti-IL-17F neutralization did not cause an increase in fungal loads in WT animals (49), suggesting that a deficiency of the IL-17AF heterodimer rather than (or in addition to) IL-17F could be responsible for increased *C. albicans* infection in *Il17f*^*S65LS66L*^ mice. A protective role of IL-17AF could also explain why IL-17A blockade failed to further promote a fungal burden in *Il17f*^*S65LS65L*^ mice. If IL-17AF is indeed the primary effector cytokine among the three IL-17RA/IL-17RC receptor ligands, the increased susceptibility caused by anti-IL-17A Ab treatment could be due to its capacity to block IL-17AF rather than the IL-17A homodimer, as generally assumed (49). Since *Il17f*^*S65L/S65L*^ mice already have a fully impaired IL-17AF signaling pathway, this could explain why anti-IL-17A antibody treatment did not lead to a higher fungal burden than isotype control antibodies.

An unexpected observation made in these studies was that the IL-17F.S65L mutation affected CXCL1 mRNA expression and subsequent neutrophil recruitment yet had no detectable impact on expression of key antifungal AMPs, *β*-defensin 3 and S100A8/A9 (calprotectin). IL-17 is a potent regulator of the neutrophil axis, acting on target epithelial cells to induce chemokines that in turn recruit myeloid cells, especially CXCR2-expressing neutrophils (62). This matters not only in OPC but other oral infections such as periodontal bone loss (95) as well as pulmonary infections (96, 97). Even so, not all studies of OPC have found that IL-17 regulates neutrophil infiltration upon OPC (98); the reason for the discrepancy is unclear, but could possibly relate to effects of different local oral microbiota (99, 100).

Although the IL-17RA/IL-17RC heterodimer is the canonical receptor complex thought to transduce signaling of IL-17A, IL-17F and IL-17AF (101, 102), recently several alternative configurations of the receptor have been proposed. IL-17RD was suggested to act in concert with IL-17RA to mediate IL-17A but not IL-17F signaling in keratinocytes to drive psoriasis-like skin inflammation (32). Thus far, the binding capacity of IL-17AF to an IL-17RA/RD receptor complex has not been characterized. In the OPC model, *Il17rc*^*-/-*^ mice phenocopy *Il17ra*^*-/-*^ mice in terms of fungal loads and other signs of disease (15), suggesting that IL-17RC is needed to mediate immunity. *Il17rc*^*-/-*^ mice are also susceptible to dermal candidiasis (103). In agreement with results in mice, humans with *IL17RC* null mutations experience CMCD (18). Hence, it is unlikely that an IL-17RC-independent receptor complex mediates host-defense during mucocutaneous candidiasis.

A recent structural analysis of the IL-17RC subunit unexpectedly revealed that IL-17F may have the ability to signal through an IL-17RC homodimeric receptor (31). Interestingly, based on this structure, the IL-17F.S65L mutation is predicted to cause steric hindrance that decreases its binding affinity to IL-17RC. However, the contacts of IL-17F to IL-17RA are less tight than to IL-17RC, and hence changes in binding affinity to this subunit would likely be less pronounced (31). Alignment of the IL-17RC residues that are proximal to IL-17F.S65 shows full conservation between the human and mouse receptors, arguing for strong overall structural conservation (*Jean Michel Rondeau, personal communication*). Putting this together, we speculate that the increased fungal burden in *Il17f*^*S65LS65L*^ mice is caused by reduced interactions of IL-17F with the IL-17RC homodimeric receptor, rather than with the IL-17RA/IL-17RC heterodimer.

In summary, results from this new IL-17F.S65L mutant mouse strain in the murine OPC model are generally, though not completely, in line with data in CMCD patients with the IL-17F.S65L mutation. The mutation leads to a reproducible, albeit mild, increase in the degree of mucosal *C. albicans* infection. However, unlike humans, the mutation does not show evidence for activity in a heterozygous configuration. These data confirm the importance of the IL-17 signaling axis in OPC. Moreover, these mice may be useful for interrogating other activities of IL-17F and possibly IL-17AF where their functions are not well defined.

## Acknowledgements

SLG was supported by NIH grants AI128991 and DE022550. DHK was supported by AR071720 and AR060744. R. Gordon was supported by T32-AI089443. We are grateful to Casey Weaver (University of Alabama) for *Il17f*^*Thy1.1*^ reporter mice and Ulrich Siebenlist (NIH) for *Act1*^*-/-*^ mice. *Il17ra*^*-/-*^ mice were a kind gift from Amgen. We thank Chunming Bi of the University of Pittsburgh Transgenic and Gene Targeting for expert technical help. Drs. Jean Michel Rondeau and Frank Kolbinger (Novartis) and Yufang Shao (Bon Opus) provided valuable discussions. We also thank Drs. Mark Shlomchik, Jean-Laurent Casanova and Anne Puel for additional input.

## Abbreviations

AMP: antimicrobial peptide;
APECED: Autoimmune polyendocrinopathy-candidiasis-ectodermal dystrophy;
CMCD: chromic mucocutaneous candidiasis disease;
ILC: innate lymphoid cell;
OPC: oropharyngeal candidiasis.

## Author contributions

Conceptualization, SLG; Methodology, LM, RG, DHK, SG; Investigation, CZ, LM, FEYA, RB, TNE; Resources, DHK; Writing – original draft, CZ, SLG; Writing – Review & Editing, CZ, LM, RG, FEYA, RB, TNE, DHK, SG, SLG; Visualization, CZ, SG, SLG; Supervision, SLG; Funding Acquisition, SLG.

---

***Supplementary Figure 1. Glycosylation site prediction in murine IL-17F***

The IL-17F amino acid sequence (see Figure 1) 1 was analyzed for predicted N-glycosylation sites (NetNGlyc 1.0 server). Asn-X-Ser/Thr sequences are shown in blue, and Asn residues predicted to be N-glycosylated are shown in red. + potential > 0.5, ++ potential > 0.5 and jury agreement (9/9) or potential > 0.75.

***Supplementary Figure 2. Genomic DNA sequencing of IL-17F.S65L mutation***

Founder mice were created by CRISPR/Cas9. Genomic DNA of founders was extracted from tail tissue and subjected to sequencing. (A) DNA sequence of mice with an *Il17f* ^*S65L/S65L*^ homozygous mutation. Black dot indicates nucleotide replacement. (B) Chromatogram of the DNA sequence data showed in panel A.

