## Supplementary material for "A loss-of-function mutation in IL-17F enhances susceptibility of mice to oropharyngeal candidiasis": Supp Figs

**Suppl. Figure 1**

```
Name:  Sequence         Length:  169
MKCTRETAMVKSLLLLMLGLAILREVAARKNPKAGVPALQKAGNCPPLEDNTVRVDIRIFNQNQGISVPREFQNRSSSPW      80
DYNITRDPHRFPSEIAEAQCRHSGCINAQGQEDSTMNSVAIQQEILVLRREPQGCSNSFRLEKMLLKVGCTCVKPIVHQA     160
AHHHHHHHH
........................................................................N.......      80
..N.............................................................................     160
.........                                                                            240

(Threshold=0.5)
----------------------------------------------------------------------
SeqName      Position  Potential   Jury    N-Glyc
                                 agreement result
----------------------------------------------------------------------
Sequence       74 NRSS   0.5910     (8/9)    +
Sequence       83 NITR   0.6987     (9/9)    ++
----------------------------------------------------------------------
```

**Suppl. Figure 2**

**A.**

| | |
|---|---|
| Mouse 1 | CGATTTCTAG AGGGAACCGG |
| Mouse 2 | CGATTTCTAG AGGGAACCGG |
| Mouse 5 | CGATTTCTAG AGGGAACCGG |
| Mouse 11 | CGATTTCTAG AGGGAACCGG |
| Mouse 13 | CGATTTCTAG AGGGAACCGG |
| Template | CGATCTCTGA GGGGAACCGG |

**B.**

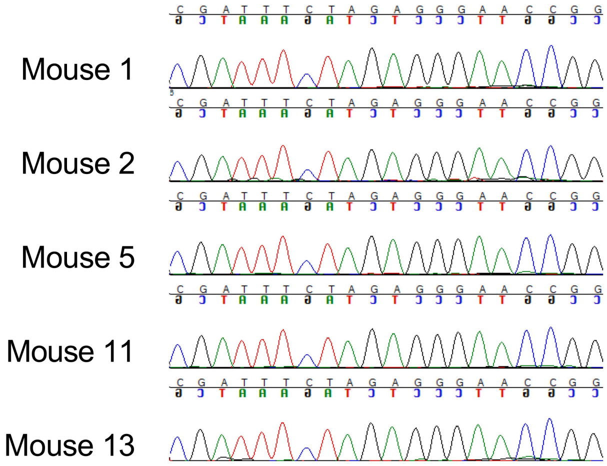
